# Global analysis of N6-methyladenosine functions and its disease association using deep learning and network-based methods

**DOI:** 10.1101/461673

**Authors:** Song-Yao Zhang, Shao-Wu Zhang, Xiaonan Fan, Jia Meng, Yidong Chen, Shoujiang Gao, Yufei Huang

## Abstract

N6-methyladenosine (m^6^A) is the most abundant methylation, existing in >25% of human mRNAs. Exciting recent discoveries indicate the close involvement of m^6^A in regulating many different aspects of mRNA metabolism and diseases like cancer. However, our current knowledge about how m^6^A levels are controlled and whether and how regulation of m^6^A levels of a specific gene can play a role in cancer and other diseases is mostly elusive. We propose in this paper a computational scheme for predicting m^6^A-regulated genes and m^6^A-associated disease, which includes Deep-m^6^A, the first model for detecting condition-specific m^6^A sites from MeRIP-Seq data with a single base resolution using deep learning and a new network-based pipeline that prioritizes functional significant m^6^A genes and its associated diseases using the Protein-Protein Interaction (PPI) and gene-disease heterogeneous networks. We applied Deep-m6A and this pipeline to 75 MeRIP-seq human samples, which produced a compact set of 709 functionally significant m^6^A-regulated genes and nine functionally enriched subnetworks. The functional enrichment analysis of these genes and networks reveal that m^6^A targets key genes of many critical biological processes including transcription, cell organization and transport, and cell proliferation and cancer-related pathways such as Wnt pathway. The m^6^A-associated disease analysis prioritized five significantly associated diseases including leukemia and renal cell carcinoma. These results demonstrate the power of our proposed computational scheme and provide new leads for understanding m^6^A regulatory functions and its roles in diseases.

**Author summary:** The goal of this work is to identify functional significant m^6^A-regulated genes and m^6^A-associated diseases from analyzing an extensive collection of MeRIP-seq data. To achieve this, we first developed Deep-m^6^A, a CNN model for single-base m^6^A prediction. To our knowledge, this is the first condition-specific single-base m^6^A site prediction model that combines mRNA sequence feature and MeRIP-Seq data. The 10-fold cross-validation and test on an independent dataset show that Deep-m^6^A outperformed two sequence-based models. We applied Deep-m^6^A followed by network-based analysis using HotNet2 and RWRH to 75 human MeRIP-Seq samples from various cells and tissue under different conditions to globally detect m^6^A-regulated genes and further predict m^6^A mediated functions and associated diseases. This is also to our knowledge the first attempt to predict m^6^A functions and associated diseases using only computational methods in a global manner on a large number of human MeRIP-Seq samples. The predicted functions and diseases show considerable consistent with those reported in the literature, which demonstrated the power of our proposed pipeline to predict potential m^6^A mediated functions and associated diseases.

## Introduction

*N*^6^-methyl-adenosine (m^6^A) methylation is a paradigm-shifting research filled with exciting discoveries. Recent research has shown that m^6^A exists in > 25% of mRNAs in mammalian cells [1, 2] and forms an important regulatory circuitry that controls many aspects of RNA metabolism [3–10]. Evidence of m^6^A’s involvement in cancer and other diseases [11–21] and its role in regulating viral life cycle [22–25] are also accumulating. However, our current knowledge about how m^6^A levels are regulated and whether and how regulation of m^6^A levels of a specific gene can play a role in cancer and other diseases is largely elusive.

The purpose of this study is to conduct a comprehensive prediction of m^6^A mediated functions and associated diseases through global analysis of m^6^A regulated genes using 75 human methylated RNA immunoprecipitation sequencing (MeRIP-seq) [1, 2] samples curated by MeT-DB2 [26]. To this end, prediction of context-specific m^6^A sites is an essential first step. Several informatics tools have been developed to predict condition independent m^6^A sites from RNA sequences [27–35] or condition-specific m^6^A peaks in MeRIP-seq data [36, 37]. Chen et al. [35] proposed the first sequence-based model iRNA-Methyl to predict m^6^A site using features extracted from RNA sequences. Subsequently, Zhou et al. developed SRAMP [29] to improve the performance of predicting single-base m^6^A sites using three kinds of features extracted from pri- and mature RNA sequences. Since then, a line of sequence-based algorithms [38–40], all based on different handcrafted features extracted from RNA sequences have been developed. As an alternative to handcrafted RNA sequence features, Wei et al. [33] proposed DeepM6APred to extract features automatically using a deep belief network (DBN). However, because RNA sequences are independent of any study conditions, none of these sequence-based models can predict condition-specific m^6^A sites. Because m^6^A has been shown to play different regulatory roles in different cell conditions and disease types, its methylation status is highly dynamic in nature, being either methylated or demethylated depending on the biological contexts. Also, m^6^A status can be experimentally manipulated through silencing or overexpressing key m^6^A related proteins, a commonly used approach for studying m^6^A functions. Therefore, predicting condition-specific single-base m^6^A site is an essential task in m^6^A research. Currently, algorithms including exomePeak [41] and MeTPeak [42] are proposed to predict context-specific m^6^A peaks from MeRIP-seq data. However, MeRIP-Seq has a limited resolution of ~100bp and its large biological and technical variations often result in high false positive rates in the predicted peaks. No existing algorithm can predict condition-specific m^6^A site at a single-base resolution.

In addition to a lack of single-base, context-specific site prediction algorithms, computational prediction of m^6^A functions has not been adequately addressed. In our previous work [43], we developed m^6^A-Driver, a network-based approach to identify m^6^A driven genes with significant functions under a specific context. However, m^6^A-Driver has several limitations. First, m^6^A-Driver to identify m^6^A driven genes in two different conditions, e.g., gene knock-down vs. normal. Therefore, m^6^A-Driver cannot be used for the intended global analysis that includes samples from multiple conditions. Second, because of the small sample size, m^6^A-Driver tends to identify a large number of significant genes, which makes it difficult to prioritize these genes. In this work, we propose to develop a new approach to address these limitations of the site or peak detection algorithms and m^6^A-Driver so that a global prediction of m^6^A mediated functions and associated diseases across samples from multiple conditions can be reliably performed.

To improve the resolution and accuracy of condition-specific m^6^A site prediction from MeRIP-seq, we propose a novel Convolutional Neural Network (CNN) [44] based method named Deep-m^6^A to detect m^6^A sites from MeRIP-Seq peak regions with a single base resolution. This model integrates mRNA sequence information with MeRIP-seq data and trained on single-base m^6^A sites identified by the miCLIP [45], the state-of-the-art high throughput technology for single-base detection of m^6^A sites. Deep-m^6^A makes it possible to identify single-base m^6^A sites from the 75 human MeRIP-seq samples with higher accuracy and resolution, providing us a chance to investigate the relationship of m^6^A methylation with gene expression and diseases in a global manner. To this end, we propose a new network-based pipeline. We hypothesize that m^6^A regulates biological processes and pathways by regulating a set of functionally interacted genes through regulating their mRNA stability. Therefore, the expression of a gene regulated by an m^6^A site is likely to be correlated with the m^6^A level across multiple conditions and this regulation also influences its neighboring genes in the PPI network. The stronger an m^6^A regulates a gene, the more significant the m^6^A-expression correlation and the higher the degree of influence on its neighbors. Our goal is to identify m^6^A regulated genes, whose expressions are correlated with their m^6^A methylations across 75 samples and which are closely interacting with other m^6^A genes in a PPI network. To achieve this goal, we adopt the HotNet2 [46] algorithm. The basic idea of HotNet2 is to diffuse “heat”, i.e., correlation of gene expression and m^6^A level in this study, of genes in a network and select the significant genes, where the “heats” they diffused to each other are high. Therefore, the m^6^A regulated genes identified by HotNet2 will in general have relatively high expression-methylation correlation and closely interacted with each other in the PPI network. However, HotNet2 still can also identify m^6^A regulated genes with lower “heat” but closely interacted with “hot” genes. After m^6^A regulated genes are identified, our pipeline prioritizes the m^6^A associated diseases by applying the random walk with restart on heterogeneous network(RWRH) method [47] on a gene-disease heterogeneous network.

We applied Deep-m^6^A and this pipeline to the 75 MeRIP-seq samples, which produced a compact set of 709 functional significant m^6^A-regulated genes and nine functional enriched subnetworks. The functional enrichment analysis of these genes and networks reveal that m^6^A targets key genes of many critical biological processes including transcription, cell organization and transport, cell proliferation and cancer-related pathways such as the Wnt pathway. The m^6^A-associated disease analysis prioritized five significantly associated diseases including leukemia and renal cell carcinoma.

## Result

### Overview of the method

The flowchart of our pipeline is illustrated in Figure 1 We intend to perform a global analysis of existing human m^6^A site in different samples to uncover m^6^A regulated functions and associated disease. To achieve this, we first set out to determine the single-base m^6^A sites. To this end, we developed a novel CNN model, called Deep-m^6^A, which takes both sequence feature and MeRIP-Seq IP reads count as an input to predict context specific single-base m^6^A sites from MeRIP-Seq data. We then applied Deep-m^6^A to predict single-base m^6^A sites for all 75 human MeRIP-Seq samples extracted from MeT-DB V2.0 [26]. Then, we extracted the m^6^A sites appeared in at least 12 samples according to a test based on Fisher’s z transformation [48] and calculated the revised Fisher’s z-transformation [49] of the Pearson correlation of methylation degree and gene expression level across all their appeared samples (see Methods and material for details). Genes that contain at least one m^6^A site that occurred in more than 12 samples and whose methylation degree and gene expression level are significantly correlated were defined as candidate m^6^A regulated genes. We selected the largest absolute z-transformed correlation from all m^6^A sites in a gene to denote the methylation-expression correlation of the gene. To further study the m^6^A regulated functions and select functional m^6^A-regulated genes, we applied HotNet2 [46] to 4 different PPI networks, taking the absolute z-transformed correlations for all candidate m^6^A-regulated genes as the heat vector, to identify significant PPI subnetworks that are regulated by m^6^A more than expected by chance. The 4 PPI networks includes BioGRID [50], HINT+HI2012 [51, 52], MultiNet [53], and iRefIndex [54]. The last three networks are used in the HotNet2 paper. The edges that are identified by all the four networks were extracted to form significant consensus subnetworks, which were then extended to include edges identified in 3, 2, and 1network, respectively (See Methods for the detail). All the genes in the extended subnetworks are defined as significant m^6^A-regulated genes and genes reported in the significant consensus networks are defined as consensus m^6^A-regulated genes. Finally, we took all these significant m^6^A-regulated genes as gene seeds and their correlated diseases according the OMIM as disease seeds and applied RWRH on a heterogeneous gene-disease network. We also built four heterogeneous gene-disease networks corresponding to the four PPI networks. For each heterogeneous gene-disease network, the top 10 ranked diseases were selected as candidate m^6^A-associated disease. A random network test was applied to calculate an empirical *p*-value for each candidate m^6^A-associated disease. The candidate disease with a *p*-value < 0.05 was selected as significant candidate m^6^A-associated disease. The significant candidate m^6^A-associated disease that identified by all the 4 networks were finally identified as m^6^A-associated disease.

**Figure 1.**
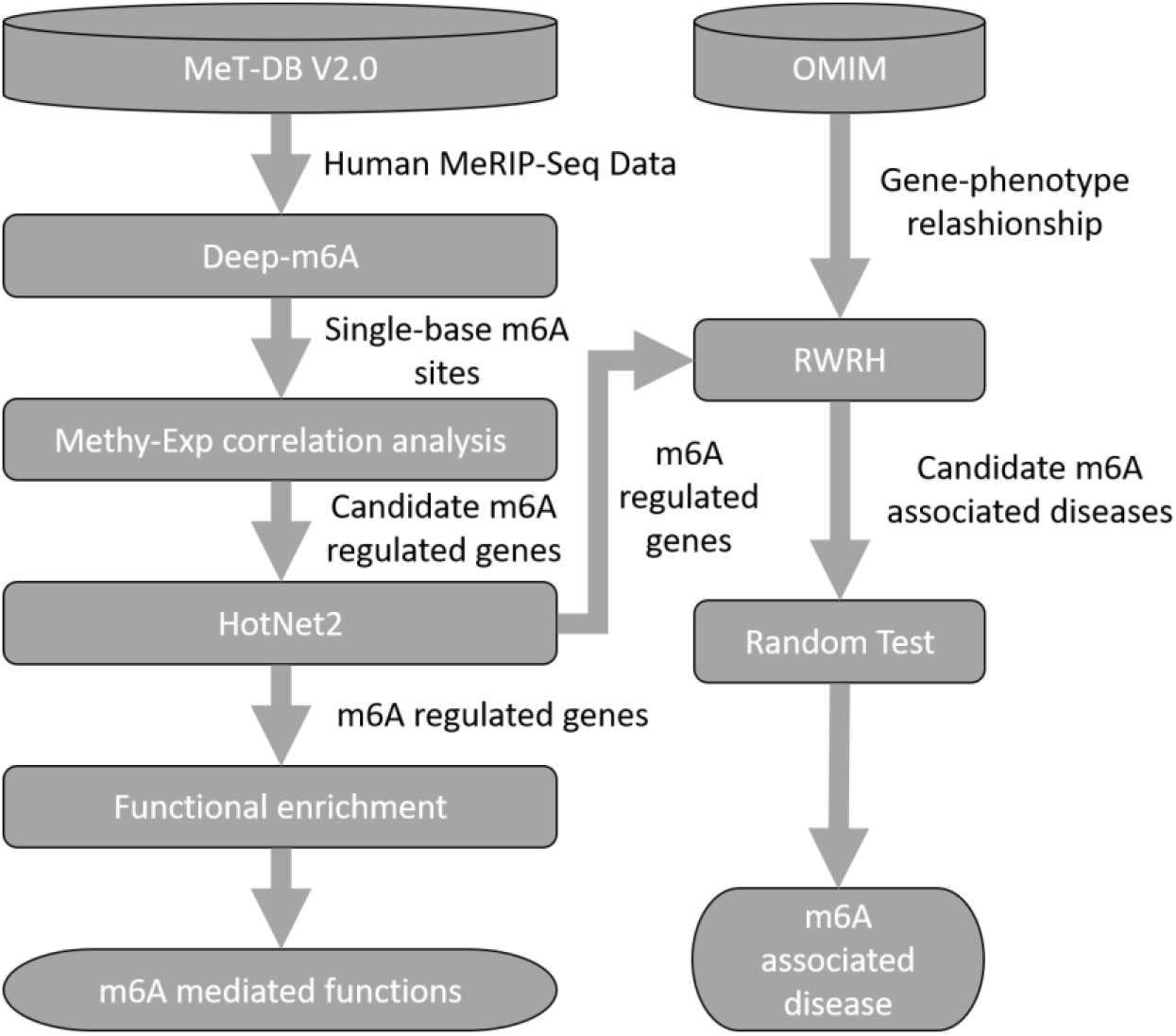
Flowchart of our proposed prediction pipeline.

### Deep-m6A achieves higher precision and sensitivity in 10-fold cross validation

We first evaluated the performance of Deep-m^6^A using the training data from the HEK293 cell lines and compared its performance with Deep-m6A-S. As described in Methods, Deep-m6A-S and Deep-m^6^A use the same CNN structure but different input features. Deep-m^6^A-S takes 101-nt long sequences centered at a DRACH motif as input, whereas Deep-m^6^A takes both sequence and the corresponding IP reads count as input. As detailed in the “Dataset” section, this training data include 4,742 positive samples that are CITS miCLIP m^6^A sites in the MeRIP-seq peak regions and also centered at DRACH motif, and 33,718 negative samples that are also centered at a DRACH motif in the MeRIP-seq peak regions but are > 50-nt away from the positive samples and not CIMS miCLIP sites or single-base m^6^A sites reported in other experiments [55]. Before model comparison, we investigated the difference between MeRIP-seq IP reads coverage in the sequence regions of the positive and negative samples. As shown in Figure 2 (A), the average reads count of positive samples are higher and more centered at the DRACH m^6^A motif than those of the negative samples.

**Figure 2.**
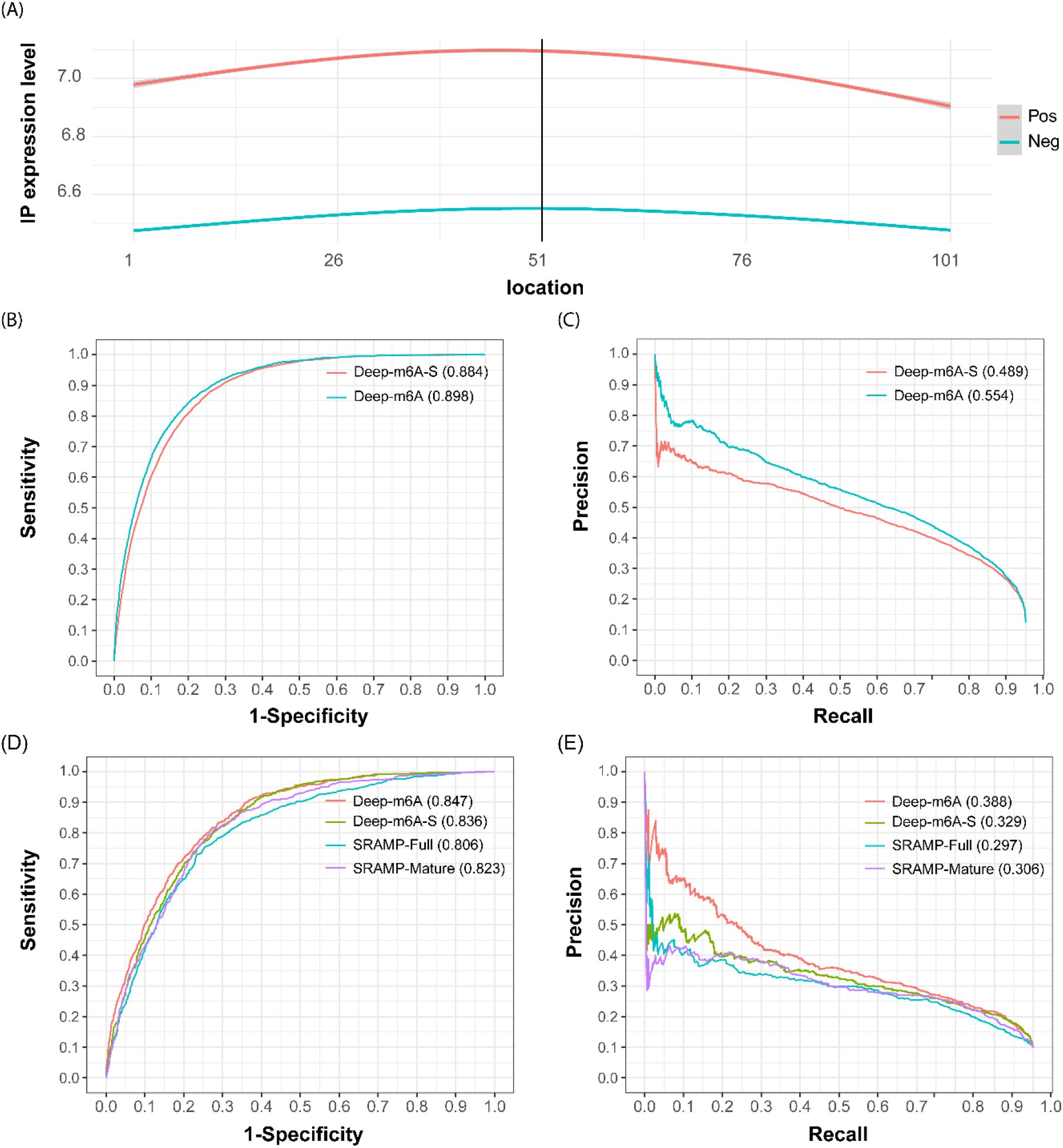
Performance of Deep-m^6^A. (A) shows the IP reads count coverage in the 101-nt positive and negative training samples. The x-axis denotes the relative location of the nucleotides in the sample sequences, where the 51st position is the center A of the DRACH motif and the 1^st^ position is the 5’ end of the sequence. The IP expression level represents the IP reads input *RC_norm_*. (B) and (C) are the ROC and PR curves of Deep-m^6^A-S and Deep-m^6^A obtained from the 10-fold CV on the HEK293 training data. (D) and (E) are performances of Deep-m^6^A, Deep-m^6^A-S, SRAMP-Mature and SRAMP-Full model on the independent MOLM13 data. The number after the method names are corresponding area under the curve (AUC).

The CNN architecture of Deep-m^6^A and Deep-m^6^A-S includes one convolutional layer followed by a max-pooling layer, a fully connected layer and an output layer. We used 10-fold cross validation to evaluate the performance of Deep-m^6^A and Deep-m^6^A-S and took one of the 10 CVs to optimize the hyperparameters. To reduce the impact of significant imbalance between the positive and negative samples, we split the negative samples into 7 subsets, each of which has the equal size to positive samples and for each CV, we trained 7 models for each balanced pair of positive/negative samples and used the average predicted score of the 7 models as the final predicted probability of a test DRACH motif to be a single base m^6^A methylation site. The CNN architecture was determined by a grid search method [56], where the searched parameters are the kernel size: 4×3, 4×4 and 4×5; # of kernels: 16, 32 and 48; the max-pooling size: 1×3, 1×4 and 1×5; # of the nodes of the fully connected layer: 8, 12, 16 and 32; the fully connected layer dropout rate: 0.2, 0.25, 0.3, and 0.5; and the softmax output layer dropout rate: 0.2, 0.25, 0.3 and 0.5. Table 1 showed the optimized hyperparameters by grid search. The categorical cross entropy loss function and Adadelta [57] optimizer were adopted in the training.

**Table 1.** Optimized hyperparameters of Deep-m^6^A and Deep-m^6^A-S.

| Hyperparameters | Kernel size | # Filters | Max-pooling size | Dropout rate1 | Node of dense | Dropout rate2 |
| --- | --- | --- | --- | --- | --- | --- |
| Value | 4x5 | 32 | 1 x 4 | 0.25 | 12 | 0.25 |

\* “Kernel size” is size of CNN kernel, “# Filters” is number of CNN filters, “Dropout rate1” is the dropout rate after max-pooling, “Node of dense” is the number of nodes in the fully connected layer after convolution layers, and “Dropout rate2” is the dropout rate after the fully connected layer.

Figure 2(B) and (C) shows the receiver operating characteristic (ROC) curves and the precision-recall (PR) curves of the CV test. We have trained seven CNN models to balance the scale of the positive and negative training samples and the predicted probability of a site is an average predicted probably of these seven models. We can see Deep-m^6^A achieved a higher area under the ROC curve (AUC = 0.898) and area under the PR curve (PRAUC = 0.554) than Deep-m^6^A-S model (AUC = 0.884, PRAUC = 0.489). The AUC and PRAUC of Deep-m^6^A are 1.4% and 5.6% higher than Deep-m^6^A-S, respectively. There is especially a higher improvement in precision. This suggests that including IP reads help lower the false positive rate, especially in the higher ranked predictions. Having a higher precision is particularly important for subsequent biological validation and functional study because in practice attention is most likely given to a limited top ranked predictions.

### Deep-m^6^A outperforms sequence-based predictions on an independent dataset

To further validate the performance of Deep-m^6^A, we applied it along with Deep-m^6^A-S on the independent MOLM13 dataset and compared their performance with SRAMP [29]. As described in the “Dataset” section, this dataset includes 726 positive and 6,577 negative samples. Deep-m^6^A and Deep-m^6^A-S were trained on HEK293 dataset as described in the previous section. For SRAMP, we downloaded the SRAMP tool from their webserver (http://www.cuilab.cn/sramp/) and applied it to the 101-nt RNA (without intron) and DNA (with intron) sequences of each sample. The RNA sequences are used by the SRAMP mature RNA model and the DNA sequences are used by SRAMP full DNA model. When feeding these sequences into the SRAMP tools, they output prob abilities of all DRACH motifs contained in the sequence. We only picked the results for the centered motif of each sample as the prediction probabilities for SRAMP. Figure 2(D) and (E) shows the ROC curve and PR curve results of these models. First, among the three sequence-based models, Deep-m6A-S reports about 1% improvement in AUC and 2% in PRAUC over the two SRAMP models. This suggests that the CNN model can capture additional discriminate sequence features than SRAMP. Second, Deep-m^6^A outperforms all three sequence-based prediction models in both AUC and PRAUC. The improvement in precision is even more pronounced (~6% over Deep-m^6^A-S). This once again speaks the benefit of including IP reads in the prediction.

### Identification of candidate m^6^A-regulated genes from all human MeRIP-Seq data

The relatively large number of human MeRIP-Seq data of different cells and tissues under different conditions curated by MeT-DB2 gives us a chance to investigate the relationship of m^6^A methylation with gene expression and diseases in a global manner. As the first step of this global investigation, we applied Deep-m^6^A to the 75 human MeRIP-seq samples to identify candidate m^6^A-regulated genes. A positive site was predicted when the prediction probability calculated by Deep-m^6^A is greater than 0.907; this threshold is chosen because at this threshold the 10-fold CV test in training can achieve a precision of 0.7. This resulted in 23,456 single-base m^6^A sites in all 75 samples. Figure 3 (A) shows the frequency of a site being predicted in the 75 samples. As shown, ~23% of these sites are sample specific, i.e. they only appear in one sample. In contrast, ~30% sites appear in more than 12 data samples.

**Figure 3.**
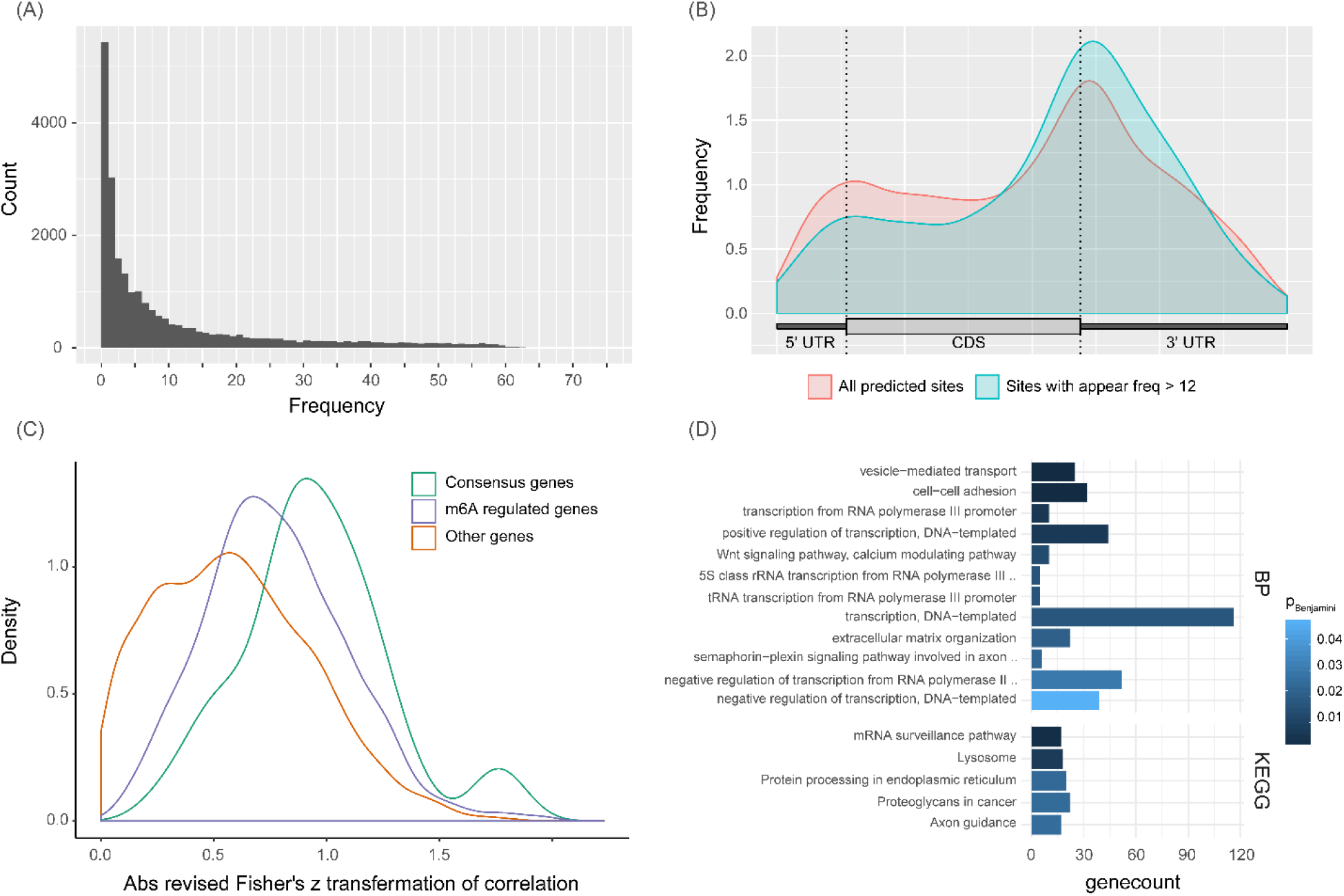
Analysis of predicted m^6^A sites in 75 human samples. (A) Distribution of the number of occurrence of a predicted single-base m^6^A site in the 75 samples. (B) Distributions of all predicted m^6^A sites and sites appeared in more than 12 samples in a meta-mRNA. (C) Distributions of the absolute revised Fisher’s z transformed Pearson correlations of the 49 consensus m^6^A-regulated genes, 709 m^6^A-regulated genes identified by HotNet2, and other remaining candidate genes. (D) GO BP and KEGG pathway enrichment result for all 709 m^6^A-regulated genes. Gene count means the number of genes involved in the corresponding terms and *P_Benjamini_* is the adjusted FDR of the enrichment p-value.

To conduct a global analysis across all the 75 samples, we extracted sites that appear in more than 12 samples. The distributions of these sites on mRNA as well as all predicted m^6^A sites were tally using Guitar R package [58] and shown in Figure 3 (B). As illustrated, for all the predicted sites, the distribution tends to be enriched in 3’ UTR and 5’ UTR, whereas the sites appeared in more than 12 samples are more enriched around stop codon and in 3’ UTR. Because 3’UTR contain binding sites of miRNA and RNA binding proteins such as HuR that are known to post-transcriptionally regulate gene expression, this distribution may indicate that the sites that appeared in >=12 samples may potentially be involved in regulating gene expression. Next, we selected the genes which harbor m^6^A sites that appear in >= 12 samples as candidate m^6^A-regulated genes. In total, 3,670 m^6^A-regulated genes that contain totally 7,090 m^6^A sites were extracted. To study the relationship of their expression and m^6^A methylation, we calculated the Pearson’s correlation of methylation degree and gene expression level for all the candidate m^6^A-regulated genes and then converted it to a revised Fisher’s z score as described in “Methods and material” section.

### Functional significant m^6^A-regulated genes identified by HotNet2 are involved in important functional pathways

We then set out to identify functional significant m^6^A-regulated genes, which are defined as candidate genes whose expression levels are influenced by the m^6^A methylation more than expected by chance and which are closely interacting with each other in PPI networks. We applied HotNet2 (See Methods and material for detailed algorithms) to the candidate m^6^A-regulated genes, taking their absolute revised Fisher’s z score as the heat vector and performing heat diffusion in the PPI network to identify functional interacted genes with relative high correlation. 49 consensus m^6^A-regulated genes were identified across all four PPI networks using HotNet2 and 709 m^6^A-regulated genes were identified in the extended network (Supplementary Figure 1; see Methods and material for detail). Most of the 709 m^6^A-regulated genes (681 or ~96%) are in the biggest subnetwork. We then compared the distributions of the absolute revised correlation z scores of the 49 consensus m^6^A-regulated genes, 709 m^6^A-regulated genes, and remaining non-significant candidate genes (Figure 3 (C)). We can see that most of the m^6^A-regulated genes have larger correlation than other nonsignificant candidate genes and the consensus m^6^A-regulated genes tend to the largest correlations among the three groups. This result speaks for the ability of our pipeline to identify likely m6A regulated genes. Notice there are some m^6^A-regulated genes with low correlations; they are identified by HotNet2 because they interacted tightly with “hot” genes with high correlation in the PPI network. To further detect the function module from these 709 m^6^A-regulated genes, we removed edges that are identified only in one PPI network, and obtained 22 subnetworks including 113 genes in total. We further isolated the 9 largest functional significant subnetworks that have at least 3 genes (Supplementary Figure 2).

We further examined the functional enrichment for all the 709 significant genes using DAVID [59] (Figure 3 (D)). Among the twelve enriched GO BPs (*P_Benjamini_* < 0.05), seven are directly related to transcription including transcription from RNA polymerase III promoter (10 genes, *P_Benjamini_* = 4.78×10^-3^), positive regulation of DNA-templated transcription (44 genes, *P_Benjamini_* = 5.79×10^-3^), transcription, DNA-templated (116 genes, *P_Benjamini_* = 1.67×10^-2^), tRNA transcription from RNA polymerase III promoter (5 genes, *P_Benjamini_* = 1.62×10^-2^), 5S class rRNA transcription from RNA polymerase III type 1 promoter (5 genes, *P_Benjamini_* = 1.62×10^-2^), negative regulation of transcription from RNA polymerase II promoter (52 genes, *P_Benjamini_* = 2.92×10^-2^) and negative regulation of transcription, DNA-templated (39 genes, *P_Benjamini_* = 4.82×10^-2^). Indeed, the m^6^A is shown to involve in every stage of RNA metabolism including transcription. It is shown to regulate mRNA stability and speculated to also regulate mRNA splicing [60]. Also, we also see that m^6^A genes could regulate cell organization and transport as they are enriched in vesicle-mediated transport (25 genes, *P_Benjamini_* = 3.83×10^-5^), cell−cell adhesion (32 genes, *P_Benjamini_* = 3.61×10^-4^) and extracellular matrix organization (22 genes, *P_Benjamini_* = 2.04×10^-2^) are also enriched. We also noticed that that the Wnt signaling pathway is enriched (10 genes, *P_Benjamini_* = 1.26×10^-2^). Wnt is an important pathway widely involved in cancer and cell development [61–63]. The potential involvement of m6A in cancer is reinforced by the enriched proteoglycans in cancer KEEG pathway (22 genes, *P_Benjamini_* = 2.43×10^-2^). This is not surprising as there is increasing evidence demonstrating its regulatory roles in different cancer [20, 64–70]. Another enriched KEGG pathways are Protein processing in the endoplasmic reticulum (20 genes, *P_Benjamini_* = 2.37×10^-2^) and Lysosome pathway (18 genes, *P_Benjamini_* = 6.31×10^-3^), both of which are protein processing pathways. These predictions are corroborated by our knowledge of the m^6^A’s role in regulating translational efficiency [71, 72].

We next performed GO and KEGG pathway enrichment analysis to the nine largest significant subnetworks (Supplementary Figure 3 & 4). These enriched BP and pathways present reasonably clear and consistent interpretations of these subnetworks. Subnetwork A is mostly involved in protein processing (6 genes enriched in Protein processing in endoplasmic reticulum, 6 genes enriched in ER-associated ubiquitin-dependent protein catabolic process, 3 genes enriched in retrograde protein transport, ER to cytosol and 4 gene enriched in protein stabilization) and potentially regulate mRNA stability via SMG9, UPF1, SMG8 and SMG1 genes enriched in nuclear-transcribed mRNA catabolic process, nonsense-mediated decay BP term. Subnetwork B is predicted to mostly involve in Notch signaling pathway via NOTCH2, MAML1 and MAML3 genes. Subnetwork C is closely related to cell motility, proliferation and survival through enrichment of m^6^A-regulated genes including VEGFB, PDGFB, VEGFA, COL5A 1, NRP1, SLIT2 and SPARC in pathways like Focal adhesion (*P_Benjamini_* = 1.75×10^-2^), positive regulation of endothelial cell proliferation (*P_Benjamini_* = 1.56×10^-3^), cell migration involved in sprouting angiogenesis (*P_Benjamini_* = 1.26×10^-3^) and positive regulation of endothelial cell migration (*P_Benjamini_* = 1.26×10^-3^). It is also involved in neuronal development related pathways including Axon guidance (NRP1, PLXNA1, SEMA3F and SLIT2 genes with *P_Benjamini_* = 1.30×10^-3^), branchiomotor neuron axon guidance (NRP1, PLXNA1 and SEMA3F genes with *PBenjamini* = 8.54×10^-3^), semaphorin-plexin signaling pathway involved in axon guidance (NRP1, PLXNA1 and SEMA3F genes with *P_Benjamini_* = 1.19×10^-3^) and axon extension involved in axon guidance (NRP1, SEMA3F and SLIT2 genes with *P_Benjamini_* = 1.19×10^-3^). The function of subnetwork D is mostly related to transcription including regulation of transcription from RNA polymerase II promoter (MED26, MED18, MED9, MED11 and MED21 genes with *P_Benjamini_* = 4.16×10^-4^), transcription, DNA-templated (MED29, MED26, POLR2I, MED9, MED11 and MED21 genes with *P_Benjamini_* = 4.92×10-3) and transcription initiation from RNA polymerase II promoter (MED26, POLR2I and MED13 genes with *P_Benjamini_* = 1.48×10^-2^). Subnetwork E’s function is defined through mostly the enrichment Wnt signaling pathway (5 genes enriched including FZD8, DKK1, LRP6, FZD5, LRP5 with *P_Benjamini_* = 6.75×10^-6^). For subnetwork F, 3 m^6^A-regulated genes are STK11, WDR6 and STRADA and they are involved in cell cycle and cell proliferation related pathways including cell cycle arrest, mTOR signaling pathway and AMPK signaling pathway with *P_Benjamini_* < 0.05. Subnetwork G serves a role in antigen processing (3 genes, TAP2, HLA-E, TAPBP enriched in Antigen processing and presentation, antigen processing and presentation of peptide antigen via MHC class I and antigen processing and presentation of endogenous peptide antigen via MHC class I with *P_Benjamini_* < 0.05). Subnetwork H includes ITGAV, ITGB5 and CYR61 as m^6^A regulated genes, which are involved in cell adhesion and subnetwork I are involved in MAPK signaling pathway via gene MAP3K4, GADD45B and MAP2K7. Taken together, these results suggest that m^6^A-regulated genes are enriched in significant biological process and pathways that can influence RNA transcription, cell motility, cell proliferation, cell survival and cell death, and therefore are likely involved in caner and cancer relate d pathways. Moreover, m6A regulated genes tend to be in the upstream of thes e enriched pathways (Supplementary Figure 5).

### Prioritized m^6^A-associated diseases

We next investigated the association of m^6^A with diseases. We first searched the OMIM database for diseases related to m^6^A-regulated genes, where we found 308 phenotypes associated with 177 m^6^A-regulated genes according to OMIM phenotype-disease relationship record (Supplementary Table 1). Most of the selected phenotypes are associated with only one m^6^A-regulated gene. To prioritize these diseases to identify significant m^6^A-associated diseases, we mapped the 709 m^6^A-regulated genes to each of the four PPI networks and the 308 phenotypes to the disease network and constructed four gene-disease networks. We then applied RWRH to each of the four networks (See “Methods and material” for the detailed algorithm), taking the regulated genes and their correlated diseases as seed nodes. Then, we selected the top 10 ranked diseases as candidate m^6^A-associated diseases and calculated an empirical *p*-value to assess if a disease association is selected by chance. Five significant m^6^A-associated diseases with *p*-value < 0.05 and reported by all the four networks were finally identified as the m^6^A-associated diseases (Table 2).

**Table 2.** Predicted m^6^A-associated diseases.

| **Disease** | **OMIM annotated m<sup>6</sup>A genes** | **Exp-m<sup>6</sup>A corr (# samples)** |
| --- | --- | --- |
| acute myeloid leukemia | NSD1, PICALM, ABL2 | 0.76 (52), 0.77 (28), 0.89 (27) |
| type 2 diabetes mellitus | HNF1B, HNF1A, WFS1, IRS2 | 0.73 (14), 0.89 (13), 0.51 (14), 0.60 (23) |
| microphthalmia | BCOR, SOX2 | 0.67 (13), 0.34 (26) |
| MHC class I deficiency | TAPBP, TAP2 | 0.81 (24), 0.46 (36) |
| renal cell carcinoma | HNF1A, OGG1, VHL, HNF1B | 0.89 (13), 0.41 (14), 0.83 (16), 0.73 (14) |

As shown in Table 2, acute myeloid leukemia is prioritized as the most significant m^6^A associated disease, where m^6^A-regulated genes NSD1, PICALM, and ABL2 are OMIM-annotated disease genes, suggesting that they may be potential m^6^A-associated biomarkers in leukemia. Several lines of evidence have shown the direct involvement of m^6^A in regulating leukemia [68–70]. The second significant disease is type 2 diabetes mellitus. Yang et al. has established this association and reported that glucose is involved in the dynamic regulation of m^6^A in patients with type 2 diabetes [73]. Our prediction identified the m^6^A genes HNF1B, HNF1A, WFS1, and IRS2 as the potential disease genes, which could provide a new clue to study the role of m^6^A in type 2 diabetes. Also, m^6^A is predicted to have a role in renal cell carcinoma. This is also corroborated by Xiao et al. [74], which reported that the m^6^A methyltransferase METTL3 acted as a tumor suppressor in renal cell carcinoma and publications in [75, 76], which identified YTHDF2, an m^6^A binding protein, to be involved in renal cell carcinoma. Taken together, we found existing evidence to support 3 out of 5 predicted association diseases. These results demonstrate the power of the proposed network-based analysis and the RWRH algorithm in identifying m^6^A associated diseases.

## Discussion

The accumulation of a large number of MeRIP-Seq samples from different cells and tissues under different conditions gives us a chance to analyze m^6^A-regulated genes and m^6^A-associated functions in a global manner. However, existing informatics tools for predicting m^6^A sites from MeRIP-seq are hampered by high false positive rates and low resolutions. To address this issue, we developed Deep-m^6^A, the first CNN model that predicts single-base m^6^A sites in MeRIP-Seq peak regions by integrating mRNA sequence features with MeRIP-Seq IP reads. Test results from 10- fold CV on training HEK293 data and an independent OMLM13 dataset showed that Deep-m^6^A outperformed sequence-based algorithms including Deep-m^6^A-S and SRAMP in both precision and sensitivity. Although the miCLIP technology has been proposed to profile transcriptome-wide m^6^A at a single-base resolution, its adoption is still very limited because of its more complex protocol. Therefore, MeRIP-seq will continue to serve as the go-to high throughput technology for global m^6^A profiling in the near future. Give the ability of Deep-m^6^A to provide MeRIP-seq a single-base detection resolution, we expect Deep-m^6^A to be an important tool in m^6^A research. Currently, Deep-m^6^A is trained to detect sites that reside on DRACH motifs. Even though this requirement of containing motifs helps reduce the false positive predictions, it also sacrifices the prediction sensitivity and will inevitably miss the positive sites that do not have any motifs. Therefore, further improvement of Deep-m^6^A in the future to be able to detect sites without motifs will provide additional value for Deep-m^6^A.

In our scheme, to further reduce the false positive rate and prioritize functional significant m^6^A-regulated genes, we examined the correlation between expression and m^6^A methylation of m^6^A-regulated genes and assessedtheir function significance using HotNet2 implemented on the PPI network. We applied Deep-m^6^A plus network-based analysis to 75 human MeRIP-seq data, which resulted in a compact collection of 709 m6A-regualted genes and severalinteracting subnetworks of m^6^A-regualted genes. Functional analysis revealed that these genes are mainly involved in transcription and Wnt pathway (Figure 2 (D)). Wnt pathway is one of the key cascades regulating cell migration and cell development, and is also tightly associated with cancer such as glioblastoma (GBM). Even though direct involvement of m^6^A in Wnt pathway has not been reported, YTHDF2, an m^6^A reader, has been shown to suppress cancer cell migration by inhibiting EMT in an m^6^A dependent manner [77]. Also, ALKBH5, an m^6^A demethylase, is reported to promote GBM tumorigenesis by stabilizing nascent FOXM1 transcripts through mediating its m^6^A levels [78] and FOXM1 has also been show to control Wnt target gene expression in GBM. It is highly likely that there are alternative pathways such as Wnt, by which m^6^A regulates cell migration and tumorigenesis. Functional enrichment of subnetworks also presents a highly consistent interpretations of their functions (Supplementary Figure 3 & 4). This suggests that the m^6^A-regulated genes in the subnetworks are likely to involve in the same processes and pathways and thus share similar functions. In terms of their enriched processes and functions, we observed again important pathways such as focal adhesion, mTOR signaling pathway and AMPK signaling pathway, which are also known to regulate cell cycle, and cell migration, and are critical in cancer.

Finally, the network-based disease association analysis on the m^6^A-regulated genes reported 5 significant associated diseases, where 3 of them have been corroborated by the existing publications. Our predictions could provide clues for the mechanisms, by which m^6^A regulating these diseases. While there is no published evidence to support the other two associated diseases, our prediction points to potentially new disease associations of m^6^A.

Last but not the least, our proposed methods have several issues that need to be further addressed in the future. First, because of the lack of miCLIP data, the model has only been trained on data from HEK293 cell line; this may not be enough to capture the features of all different kinds of true positive m^6^A sites. Second, the scale of phenotype-gene relationships in the OMIM database are relative small, which might not be able to capture all potential significant disease-gene correlations. A potential solution is to integrate other disease-gene annotation information like DisGeNET [79] to make the network more complete. Finally, we only analyzed the common m^6^A sites across many samples and their influence on gene expression. However, some of these context specific m^6^A sites appear only in a unique sample and they could be important for understanding m^6^A functions under this specific condition. Developing algorithms to detect these unique genes would be another future work.

## Methods and materials

### Datasets

The positive single-base m^6^A sites for our training data for Deep-m^6^A and Deep-m^6^A-S were obtained from [45], which developed m^6^A individual-nucleotide-resolution cross-linking and immunoprecipitation (miCLIP) technology. This paper includes two alternative technologies including cross-linking–induced mutation sites (CIMSs) and cross-linking–induced truncation sites (CITSs) for human embryonic kidney (HEK293) cells using total cellular RNA. The CIMS miCLIP uses Abcam as antibody and C→T transitions as feature, whereas the CITS miCLIP uses SySy as antibody and truncations as feature. We chose CITS miCLIP generated single base m^6^A sites as true positive m^6^A sites because the corresponding MeRIP-Seq data of the HEK293 cell line that we used for training were also generated with SySy as antibody [77]. Another reason is that 74% (4847/6543) of CITS sites can be mapped to peaks generated by MeRIP-Seq data, whereas this ratio is only 55% (5202/9536) for CIMS sites. The MeRIP-Seq peaks were detected using the exomePeak R package [41] from 2 replicates of MeRIP-Seq samples [77]. A consistent peak that appears in both replicates and also contains at least one CITS miCLIP site was determined as a single-base m6A-containing MeRIP-Seq peak. The mRNA sequences of these peaks (without introns) were subsequently extracted and any sites in the sequences that contain DRACH motifs were defined as candidate m^6^A sites. A candidate m^6^A site was then extended to 101 nt centered at the “A” of the DRACH motif to capture the sequence and reads count features around the motif. The candidate sites that also are CITS miCLIP m^6^A sites were determined as the positive samples (4,742 in total) and other candidate sites that are at least 50-nt away from a positive “A” and are not any CIMS miCLIP sites or single base m^6^A sites reported in other experiments [55] were defined as the negative samples (33,718 in total).

The independent miCLIP test data were obtained from [69], which has 3 miCLIP replicates for the leukemia cell line MOLM13, each of which contains 8,113, 2,886 and 2,050 miCLIP sites, respectively. There were in total 11,746 miCLIP m^6^A sites after combining all the three replicates, whereas only 136 sites were common across all the three replicates. Here, we considered miCLIP m^6^A sites that appeared in at least two replicates as true positive m^6^A sites (1,147 in total). The corresponding MeRIP-Seq test data were obtained from [80], which included 2 MeRIP-Seq replicates for the MOLM13 cell line. One of the 2 replicates has very low sequencing depth, so we took the combined peaks of these two replicates reported by exomePeak as MeRIP-seq peaks. Similar as the training data, the MeRIP-Seq peaks that contain at least one miCLIP m^6^A sites were determined as single-base m^6^A-containing MeRIP-Seq peaks and the sites with DRACH motifs in these regions were defined as candidate single-base m^6^A sites. The candidate m^6^A sites that are also miCLIP m^6^A sites were determined as positive samples (726 in total) and other candidate sites that are >50-nt away from any positive sites and are not miCLIP sites from any of the three miCLIP replicates were determined as negative samples (6,577 in total).

The human MeRIP-Seq data were downloaded from MeT-DB V2.0 [26], which include 75 human samples from different human cell lines and tissues under different conditions, including 9 cell lines (A549, Dendritic cells, embryonic stem cell, Endoderm, HEK293T, HeLa, HepG2, neural progenitor cells, OKMS inducible fibroblasts and U2OS) and brain tissue samples under different conditions. The reference PPI networks were built based on BioGRID (release 3.4.128) [50], HINT+HI2012 [51, 52], MultiNet [53], and iRefIndex [54]. After removing the isolated proteins and self-interaction proteins, we established a PPI network with a total of 16,062 proteins and 152,676 interactions. The last three PPI networks were downloaded from http://compbio.cs.brown.edu/pancancer/hotnet2/. The HINT+HI2012 network contains 9,858 genes and 40,704 edges; the iRefIndex network contains 12,128 genes and 91,808 edges; and the Multinet network contains 14,398 genes and 109,569 edges. Diseases ontology terms were collected from the Disease ontology [81]. The “doSim” function of R package “DOSE” [82] was used to calculate the semantic similarity between two DO terms, which are used as the edge weight of the disease network. The disease gene relationship information was extracted from the OMIM database [83]. The OMIM ID was mapped to DO ID so that we can use OMIM gene–phenotype relationship to connect the PPI network and DO disease network to construct a gene-disease network. Four gene-disease heterogeneous networks were built for each of the PPI networks.

### Deep-m6A and Deep-m^6^A-S for single-base m^6^A prediction

We developed 2 CNN models for single-base m^6^A prediction, one called Deep-m^6^A and the other was Deep-m^6^A-S. These 2 models use the same CNN structure and hyperparameters but with different inputs. For Deep-m^6^A-S, the input is the OneHot encoded 101nt RNA sequences centered at the “A” of a DRACH motif. OneHot encoding translates the A, U, C, G characters into a binary vector of (1,0,0,0), (0,1,0,0), (0,0,1,0) and (0,0,0,1), respectively. Therefore, the input of Deep-m^6^A-S becomes a 4 x 101 matrix, *M_s_*. On the contrary, Deep-m^6^A takes both RNA sequences and the features of MeRIP-Seq IP reads count at each nucleotide of the RNA sequence. However, the input for Deep-m^6^A is similar to OneHot encoded *M_s_* for Deep-m^6^A-S but with 1s replaced by the IP reads count feature at that nucleotide. The IP reads count features were calculated by the same approach as exomePeak, i.e.,

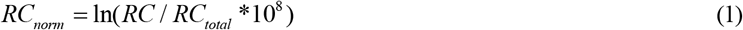

where *RC_norm_* is a 101-dimension vector of normalized reads count, *RC* is another 101-dimension vector of raw reads count, and *RC_total_* is the total number of reads for the MeRIP-Seq IP sample. Because there are two replicates in each of the HEK293 and MOLM13 datasets, we calculated the average *RC_norm_* of the two replicates as input IP reads count. The Deep-m^6^A input is then calculated as

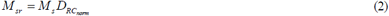

where *D_RCnorm_*is a 101×101 diagonal matrix with diagonal entries being those of *R_Cnorm_*.

Deep-m^6^A and Deep-m^6^A-S use CNN [44] to capture the non-linear features of input sequences and IP reads count. The adopted CNN architecture consists of a convolutional, a max-pooling and two fully connected layers (Figure 4). The convolutional layer outputs the pointwise product between the input matrix (*M_sr_* for Deep-m^6^A and *M_s_* for Deep-m^6^A-S) and filters, which is followed by a rectified linear (ReLU) activation. Then, a max-pooling layer, which selects the maximum value over a window, is applied to reduce the dimensionality, which is followed by a dropout operation to reduce the complexity of the model. Finally, a fully connected dense layer is added followed by another ReLU and dropout to combine all the features learned by each filter. The output of the dense layer is passed on to the softmax function to generate the probability of the input sample to be an m^6^A site.

**Figure 4.**
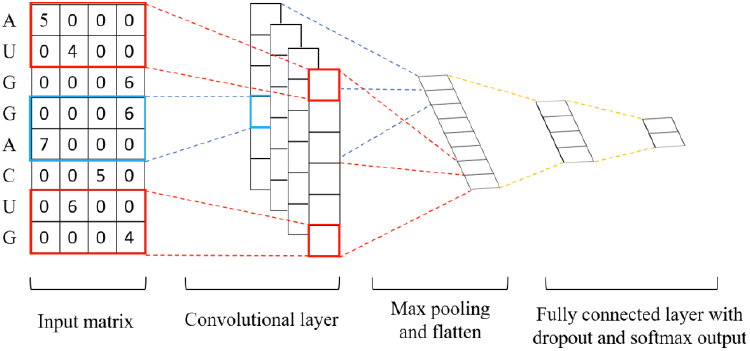
The architecture of the proposed CNN model for Deep-m^6^A. The input matrix is *M_sr_* for Deep-m^6^A and *Ms* for Deep-m^6^A-S.

Because the positive and negative sample sizes are imbalanced, we split the negative samples into seven subsets of equal size and trained 7 CNN models, each on a set that paired positive samples with a subset of negative samples. Seven models are trained as a result. For any prediction, the averaged prediction probability of the seven models is taken as the final predicted probability for an input sample. We implemented the CNN models using the keras R package. We refer Deep-m^6^A and Deep-m^6^A-S as the combination of the seven models. Input data files and Deep-m^6^A source code and the models are made publicly available on GitHub (https://github.com/NWPU-903PR/Deepm6A.git). The inputs of the Deep-m^6^A function are a MeRIP-Seq IP sample bam file and a bed file that annotates the peaks identified by exomePeak R package from MeRIP-Seq data. The output is an excel file, where each row contains information about a predicted single-base m^6^A site extracted from an exomePeak peak region and each column denotes the chromatin, the chromatin start, the chromatin end, Entrez gene ID, the predicted probability, the strand and the motif at this location, respectively.

### Prediction of single base m^6^A peak for all human MeRIP-Seq data

We applied Deep-m^6^A to predict single-base m^6^A sites in 75 human MeRIP-Seq samples from MeT-DB2. First, we applied exomePeak to predict m^6^A peaks in each MeRIP-Seq sample and then searched for DRACH motifs in the peak regions. “A”s in these motifs were treated as candidate single-base m^6^A sites and the 101nt RNA sequences centered at these “A”s and the corresponding IP reads counts were extracted to construct the input matrix *M_sr_*. For each candidate site, we separately applied Deep-m^6^A trained on the training data to calculate the probability of it to be a m^6^A site. We predict a positive site when the output probability is greater than 0.907; this threshold is chosen because at this threshold the 10-fold CV test in training can achieve a precision of 0.7. In this way, we detected in total 23,456 single-base m^6^A sites.

### Identification of m^6^A-regulated genes

We define m^6^A-regulated genes as genes whose expression level is influenced by its m^6^A methylation level more than expected by chance. To determine genes whose expression levels are significantly influenced by m^6^A, we assess the correlation between the methylation level and the corres ponding expression. If the expression levels of a gene change together with its m^6^A levels across more samples, then there will be a higher chance that the gene expression is influenced by m^6^A. To obtain a statistically meaningful correlation, we need to have the same m^6^A sites appearing in many samples. In this study, we estimated the sample size needed to detect a correlation of 0.8 with a significance level of α = 0.05. Using a two-sided test based on the Fisher’s z transformation, at α = 0.05 with power 90%, the required sample size is approximately 12 (i.e., n=12) [48]. Thus, we select single-base m^6^A sites that appear in at least 12 samples and calculated the Pearson’s correlations of the methylation degree and gene expression level across all occurred samples. The methylation degree was calculated as:

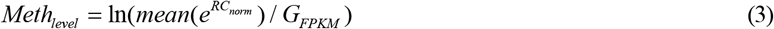

where *R_Cnorm_* is defined in (1), *G_FPKM_* is the FPKM of gene harbored the corresponding m^6^A site, calculated from the MeRIP-seq input sample by Cufflinks [84]. The expression level is denoted by the *ln* scale gene FPKM. The correlation coefficient is then transformed into a z score using a revised Fisher’s z-transformation (which is known as Hotelling’s (1953) second-order transformations) [49] that considers the influence of sample size *n*, which is expressed as

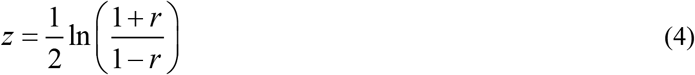

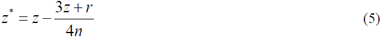

where *r* is the correlation, *n* is the sample size, and *z* is Fisher’s transformation. We used *z*\* of each gene as its correlation for the subsequent analysis in the pipeline. Genes containing this kind of m^6^A sites are treated as candidate m^6^A-regulated genes. For a candidate gene that has multiple m^6^A sites, the *z*\* score with the largest absolute value among all its harbored m^6^A sites is selected as the correlation degree for that gene and the absolute *z*\* score are used to denote the degree of regulation.

To further select m^6^A regulated genes more than expected by chance, we applied HotNet2 [85], which was originally developed to identify significant mutation genes for cancer. HotNet2 takes the gene mutation frequency or mutation score as input heat vector and applies a heat diffusion method to identify subnetworks of a genome-scale interaction network that is mutated more than expected by chance. The advantage of HotNet2 is that it not only detects significant mutated genes with high mutation frequency but also can identify significant mutation genes with relatively low mutation frequency but interact closely with other significant genes. In our case, we also want to identify significant m^6^A-regulated genes that not only have high m^6^A-expression correlations but also cooperate with each other functionally in a functional network. To apply HotNet2, we assigned absolute m^6^A-expression correlation coefficients *z*\* score of candidate m^6^A regulated genes as the input heat vector and choose the reference functional network as the four PPI networks, where BioGRID has been shown to contain the significant interactions between m^6^A methylated genes [43] and the other three PPI networks were used in the HotNet2 paper.

HotNet2 has two parameters *β* and *δ*; *β* is the fraction of the heat that a node in the network retains for itself at each time step and *δ* is the threshold, which determines whether there is an edge between 2 nodes in the final subnetwork. *β* can influence the amount of heat that a gene shares with its neighbors and is determined by the topology of the PPI network. For the three networks used in the HotNet2 paper, we selected *β* as 0.4 for HINT+HI2012, 0.45 for iRefIndex, and 0.5 for Multinet, which are reported values in [46]. For the BioGRID network, we performed the same analysis as the HotNet2 paper did to determine the value of *β*. For each different *β* from {0.05, 0.1, 0.15,…, 0.95}, we analyzed the inflection point at which the heat kept in the direct interacting neighbors of a gene drops and selected *β* as the one with the largest inflection point. As shown in Supplementary Figure 6, *β* is selected as 0.5 for the BioGRID PPI network. On the other hand, *δ* influences the scale of the subnetwork that we generated. To automatically determine it, we also performed the same analysis as did in the HotNet2 paper. For each PPI networks, we firstly generated 100 random PPI networks [86]. We take the same “heat” vector to run HotNet2 and select *δ* as the minimum one where all strongly connected subnetwork components identified by HotNet2 have a size less than and equal to a threshold *L_max_*. For each *L_max_* = 5, 10, 15, 20, we reported the median of the 100 *δ_min_*_s_ for the 100 random networks. For each run, we selected the smallest *δ* with the most significant (P < 0.05) subnetwork sizes *k* as described in HotNet2 [46]. The P-value, which denotes the significance of subnetwork size *k* is computed for the statistic *X_k_*, the number of subnetworks of size ≥ *k* reported by HotNet2. To compute an empirical distribution of *X_k_* for computing the P-value, we permuted the heat scores, i.e., the z transformation of correlation, among the genes in the original PPI network for 1000 times and then applied HotNet2 to the network using the permutated heat scores. In the end, *δ* was selected as 0.00769 for BioGRID, 0.0148 for HINT+HI2012, 0.00998 for iRefIndex, and 0.00777 for Multinet.

We applied HotNot2 to each of the four PPI networks and pooled all the genes and edges reported in at least one network to form a candidate m^6^A regulated gene network, G. We then assigned the number of the PPI networks in which an edge exists as the weight of this edge. Next, we initialized the consensus subnetworks C as the connected components of G with edges of the weight = 4, i.e., the edges that are reported in all 4 PPI networks. Then we extended each consensus subnetwork S ∈ C by adding edges of weight = 3, and afterward further extended them by adding edges of weight = 2 and then 1. Finally, we defined the genes in the final extended networks as the m^6^A regulated genes and the genes in the initial consensus subnetworks as the consensus m^6^A regulated genes. To further detect the functional modules of m^6^A regulated genes, we removed edges with weight = 1 in the m^6^A regulated subnetworks to generate functional significant subnetworks.

### Prioritize m^6^A-associated diseases

To further reveal the association between m^6^A and disease, we adopted the RWRH (random walk with restart on heterogeneous network) [47] algorithm to prioritize the candidate m^6^A regulated diseases. RWRH is a well-known heterogeneous network-based algorithm to infer the gene-phenotype relationship. The heterogeneous network contains three parts: gene-gene interaction network, phenotype (disease) network, and gene-phenotype relationship. Similarly, we also built four heterogeneous networks based on each of the four PPI networks. The disease network was generated from the semantic similarities between two DO terms of Disease Ontology database and gene-phenotype relationships were extracted from OMIM. Specifically, the OMIM phenotype IDwas mapped to DO disease ID to match the disease network with the gene-phenotype relationship. RWRH performs a random walk with restart from certain gene and disease seed nodes in the heterogeneous network. After several steps of random walk, the probability of each node, that the seed genes and seed diseases will walk to, become steady and is returned as vectors 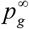 and 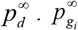 denotes the probability that the seed genes and seed diseases will walk to gene *i* and 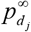 is the probability that the seed genes and seed diseases will walk to disease *j*. In another world, RWRH can prioritize how closely a gene or a disease is correlated with the seed genes and seed diseases.

To study the correlation between m^6^A and diseases, we first mapped all the 709 significant m^6^A-regulated genes obtained from HotNet2 to OMIM gene-phenotype relationship, which resulted in 308 phenotypes regulated by 177 m^6^A genes. Most of these phenotypes contain only one disease gene(The details of the m^6^A gene-disease relationships are included in Supplementary Table 1). Next, to obtain candidate m^6^A-associated diseases, we mapped these 308 m^6^A gene-correlated OMIM phenotypes to DO ID, which resulted in 90 m^6^A gene-associated DO diseases.

For each of the four heterogeneous networks, these 90 diseases were taken as disease seeds for RWRH, whereas all the 709 significant m^6^A-regulated genes were mapped to the PPI network to serve as gene seeds for RWRH. Finally, we applied RWRH using these genes and disease as seed nodes to prioritize all the DO diseases in the heterogeneous networks. Figure 5 illustrates the heterogeneous network in this study. As shown, there is a chance that *d*_3_ may be ranked as significant m^6^A-associated diseases even though there is no m^6^A-regulated gene directly connected with it. The reason is that the genes (i.e., *g*_3_, *g*_7_) that closely interacted with m^6^A regulated genes (i.e., *g*_5_) are connected to *d*_3_ and m^6^A regulated genes (i.e., *g*_5_) connected with disease (i.e., *d*_2_, *d*_4_) also closely interacted with *d*_3_. RWRH can efficiently capture this significant network topology and prioritized *d*_3_ as significant, which showed the power of this study to prioritize the potential m^6^A-associated disease with no prior knowledge that correlated with m^6^A-regulated genes.

**Figure 5.**
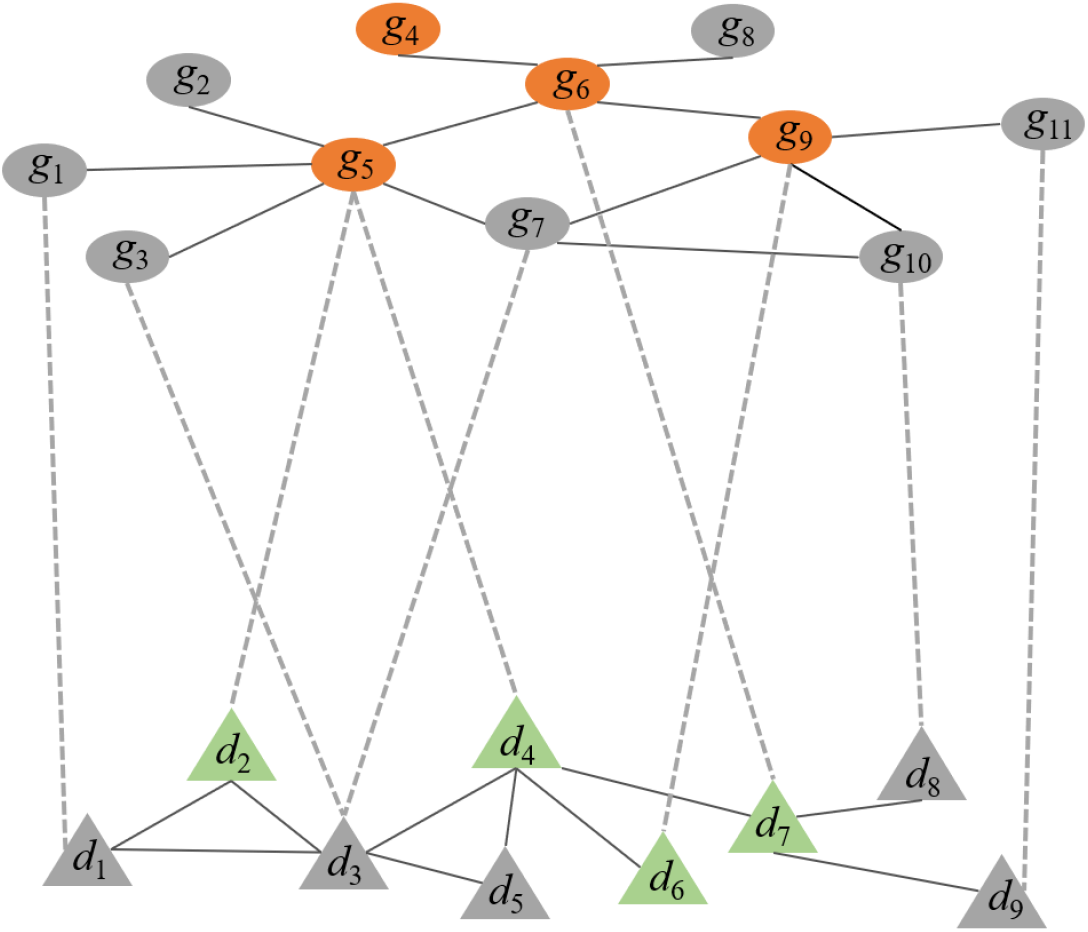
Gene-disease heterogeneous network. The top network is gene-gene interaction network, the bottom network is disease-disease similarity network and they are connected by gene-disease relationship (dashed grey lines). The orange gene nodes denoted the m^6^A regulated genes and the green disease nodes are m^6^A regulated genes correlated diseases.

The top 10 ranked diseases based on the RWRH output probability 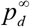 were selected as candidate m^6^A-associated diseases. To ensure these top diseases were really influenced by m^6^A-regulated genes rather than by chance, we implemented a random test to calculate an empirical p-value to assess if the disease is randomly selected. For this test, 100 random PPI networks that only keep the degree distribution of original network were generated using the method in [86]. Then, for each random PPI network, we connected it with the disease network using the disease-gene relationship and deleted the relationship between m^6^A-regulated genes and their corresponding diseases to generate a corresponding random heterogeneous network whose gene interaction relationship is random and contains no prior knowledge of the relationship between m^6^A regulated genes and any disease. After that, we applied RWRH in each of the 100 random heterogeneous networks. Finally, for each candidate m^6^A-associated disease *d_j_* (*j* = 1, 2, ⋯, 10), the empirical p-value was calculated as: 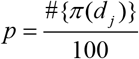, where π(*d_j_*) is a random heterogeneous network in which *d_j_* is found as the top 10 candidate disease after RWRH. The empirical p-value is calculated as the probability that a candidate m^6^A-associated disease is selected by random. The candidate m^6^A-associated diseases that with *p* < 0.05 were selected as significant candidate m^6^A-associated diseases. The significant candidate m^6^A-associated diseases that reported by all the four heterogeneous networks are determined as significant m^6^A-associated diseases.

## Acknowledgement

This work was supported by the National Natural Science Foundation of China (http://www.nsfc.gov.cn/ 61473232 and 91430111) awarded to SWZ; and the National Institutes of Health (https://www.nih.gov/R01GM113245) awarded to YH. We thank the computational support from UTSA’s HPC cluster Shamu, operated by the Office of Information Technology.

## Supporting Information

**S1 Figure.**
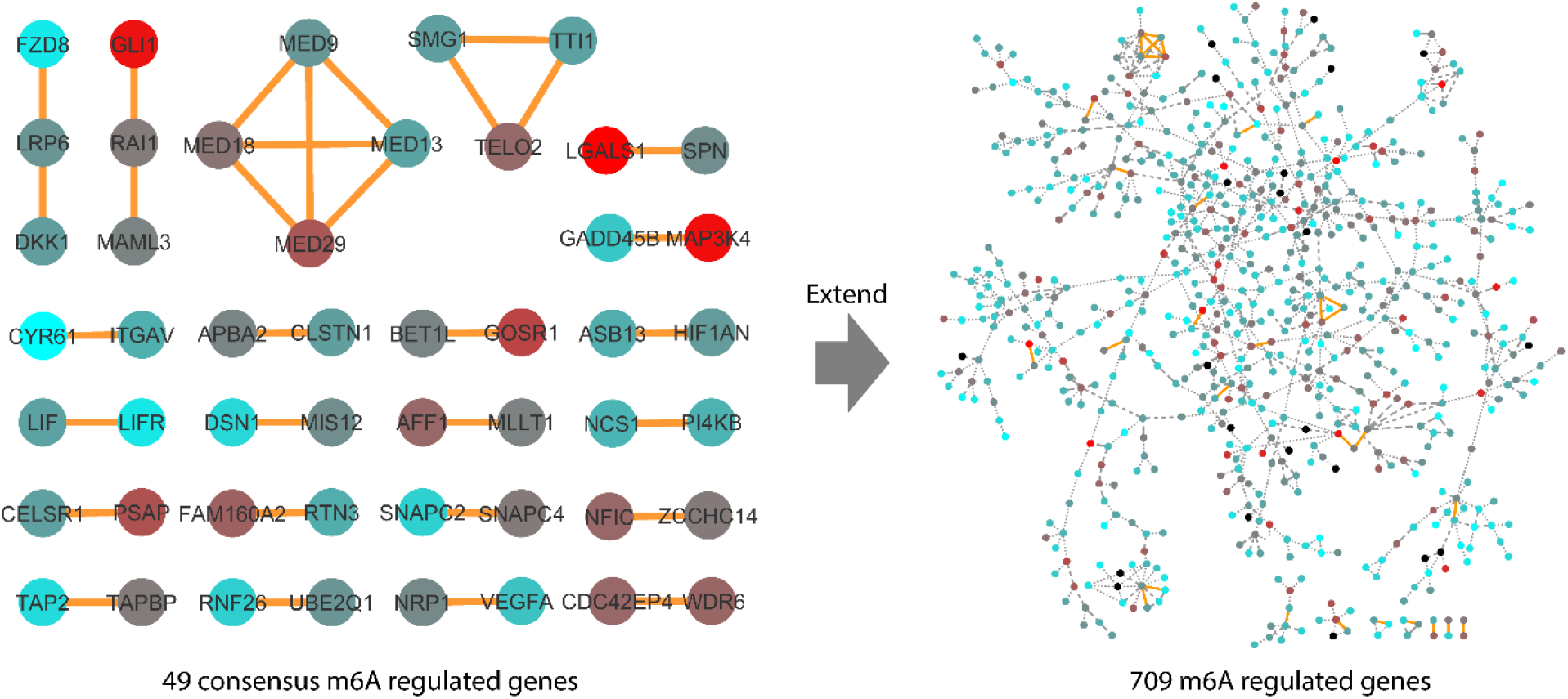
Consensus m^6^A-regulated subnetworks and m^6^A-regulated subnetworks. The color of the node denotes the heat i.e. expression-methylation correlation degree it has, red denotes higher and blue denotes lower. The solid orange edge is edge that identified in all 4 PPI networks, the dashed edge is edge that identified in at least 3 networks, the dash dot edge is in 2 networks and the dotted is 1.

**S2 Figure.**
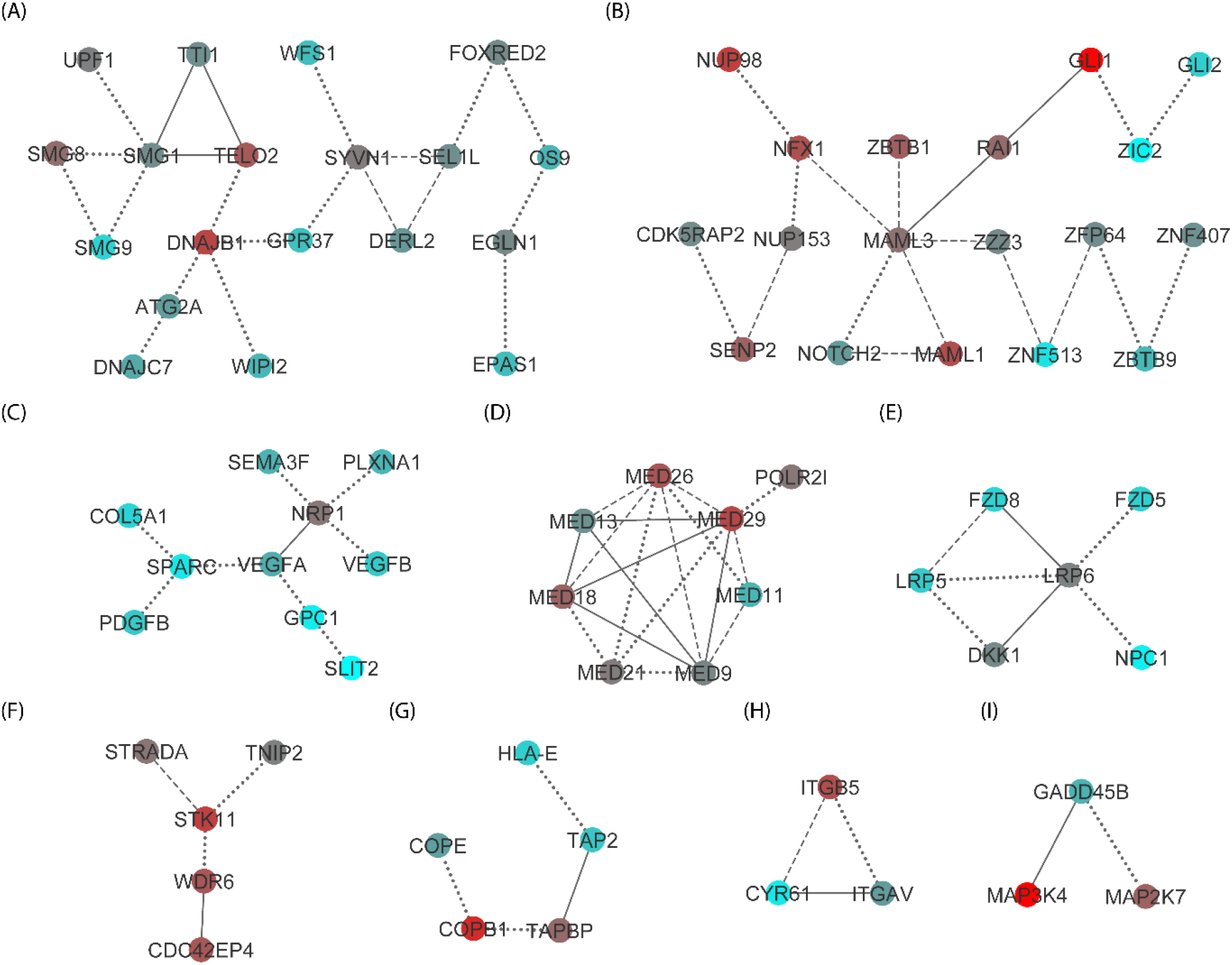
Functional significant m^6^A-regulated subnetworks. The color of the node denotes the heat i.e. expression - methylation correlation degree it has, red denotes higher and blue denotes lower. The solid edge is edge that identified in all 4 PPI networks, the dashed edge is edge that identified in at least 3 networks and the dotted edge is in 2 networks.

**S3 Figure.**
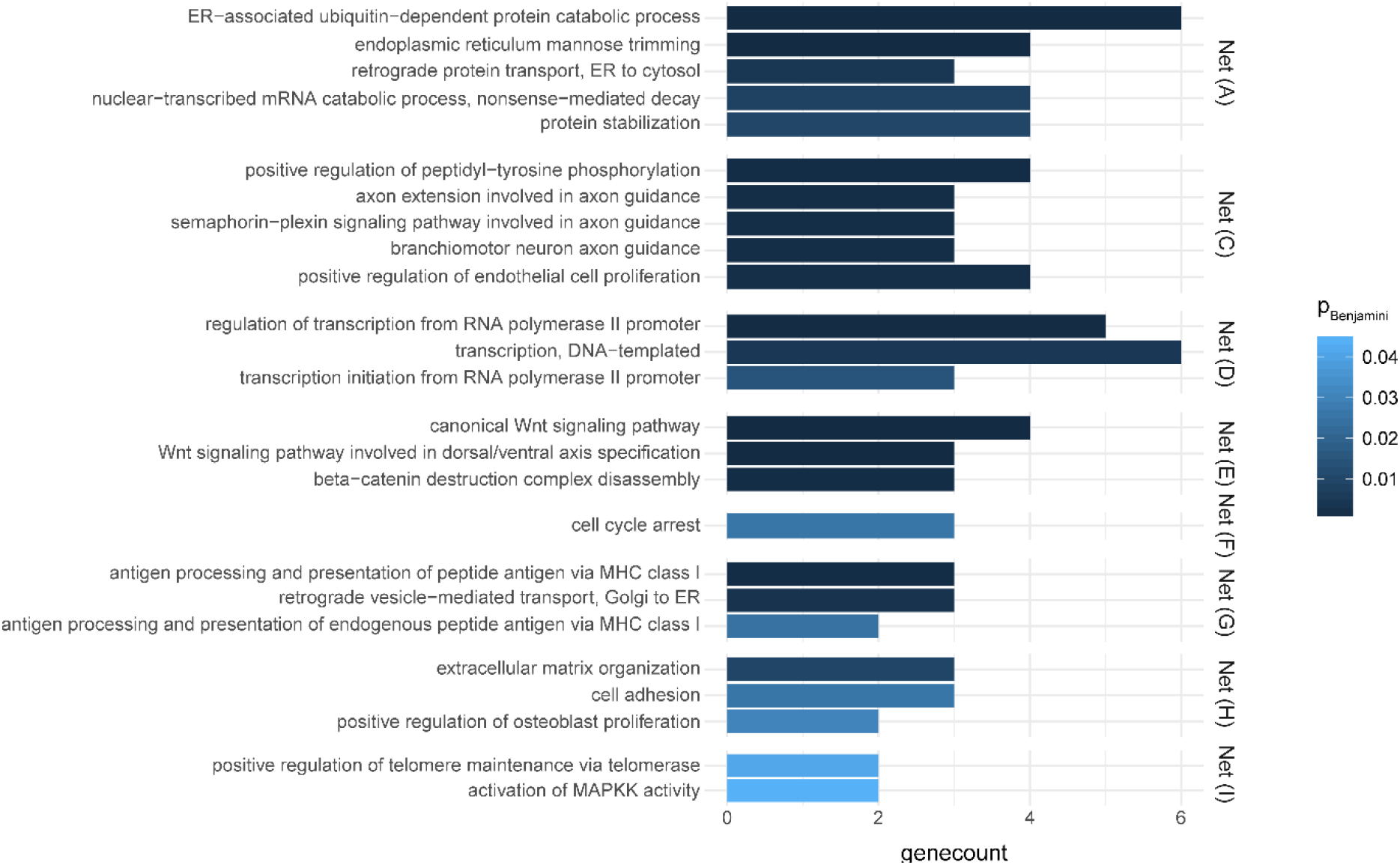
Significantly enriched GO BP terms for each of the 9 largest significant subnetworks. There is no significant BP for subnetwork (B). Gene count means number of genes involved in the corresponding BP terms and *P_Benjamini_* is the adjusted FDR of enriched p-value. All the involved BP terms have a *P_Benjamini_* < 0.05 and we only list the top 5 if the enriched terms are more than 5.

**S4 Figure.**
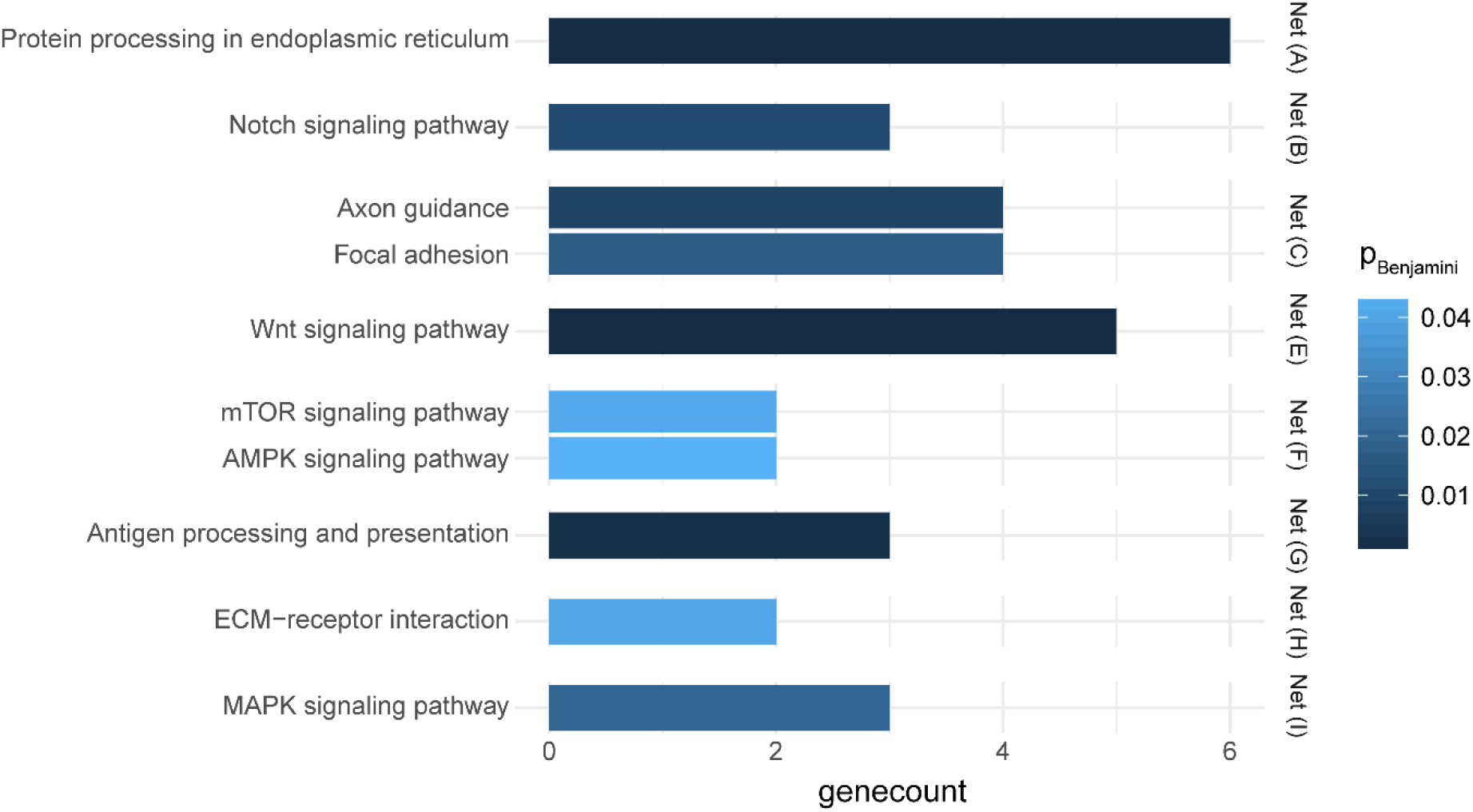
Significantly enriched KEGG pathways for the 9 largest significant subnetworks. There is no significant KEGG pathways for network (D). Gene count means number of genes involved in the corresponding pathways and *P_Benjamini_* is the adjusted FDR of enriched *p*-value. All the involved pathways have a *P_Benjamini_* < 0.05.

**S5 Figure.**
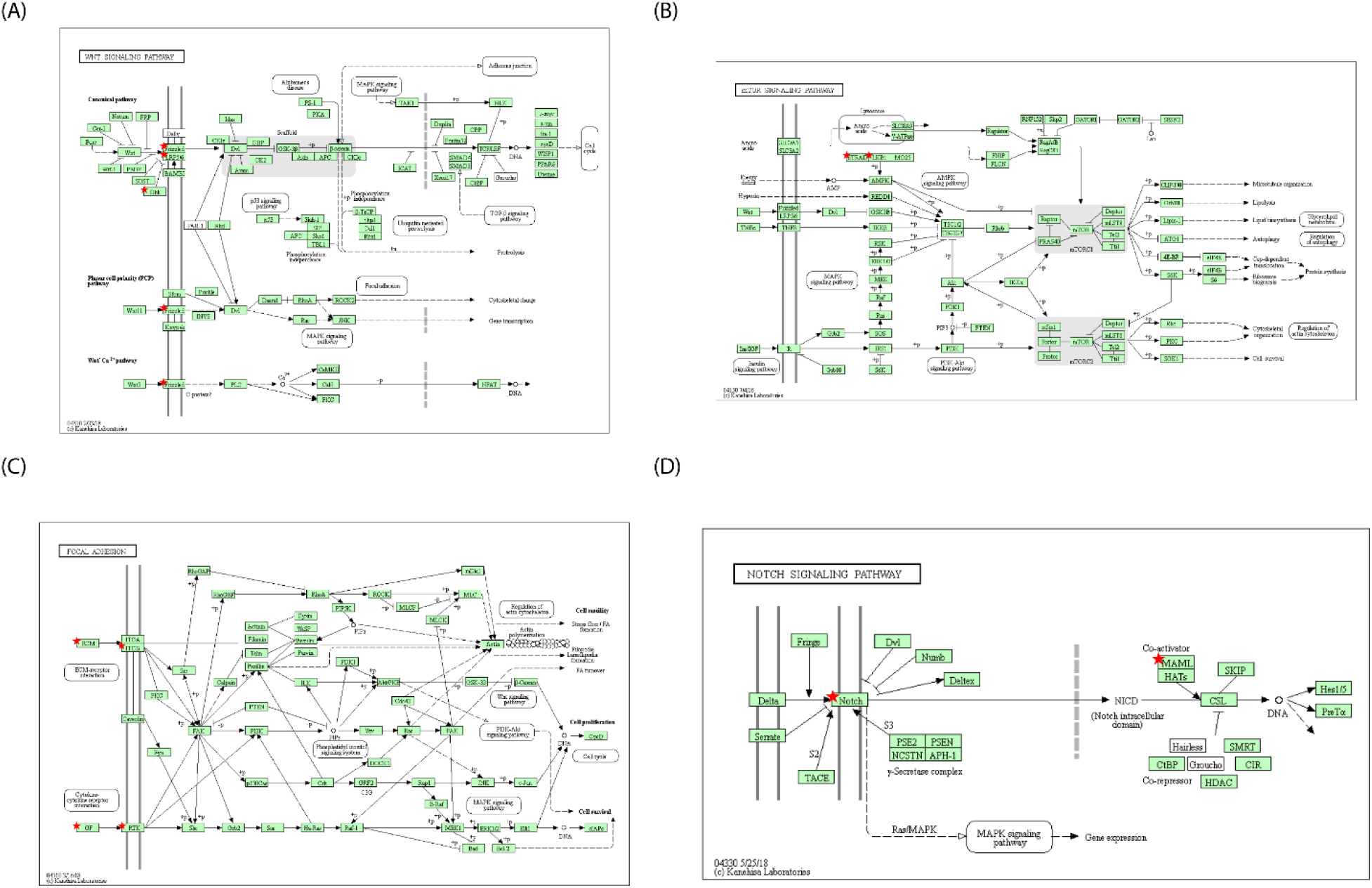
Parts of the enriched KEGG pathways of the 9 significant subnetworks. The genes marked with red star are m6A regulated genes in the subnetworks. As is shown, m^6^A regulated genes tend to be in the upstream of these enriched pathways.

**S6 Figure.**
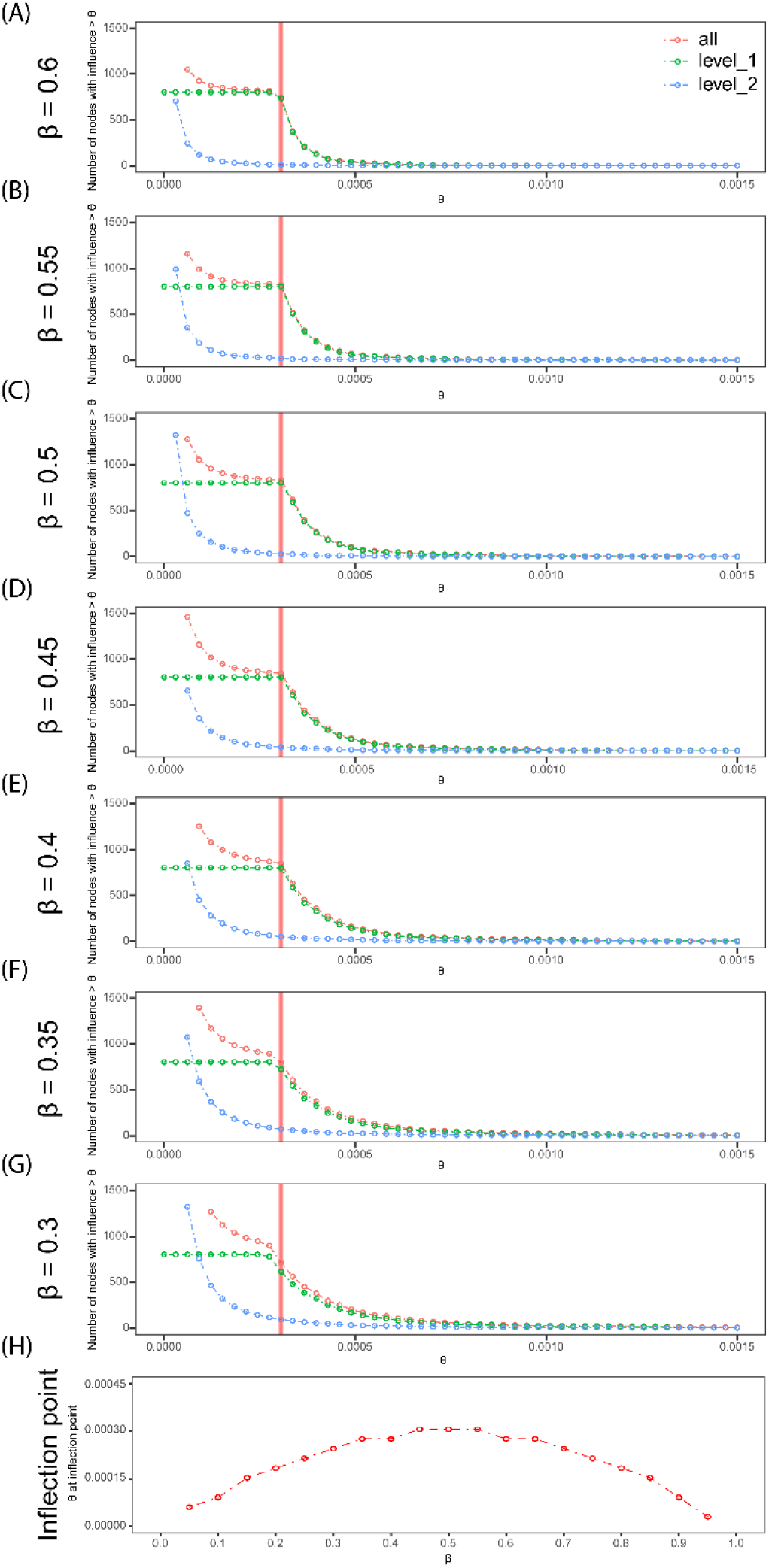
Selection of *β* of the BioGRID network. Figures (A)-(G) represent the distributions of number of nodes with influence larger than a cutoff for *β* from 0.6 to 0.3 for gene TP53, which has a high betweenness centrality. The *x*-axis of each distribution represents *θ*, a cutoff of influence. The *y*-axis represents the number of nodes in the interaction network with influence larger than *θ*. Red dotted circles denote the number of all nodes with an influence larger than different *θ*s, green dotted circles denote that of the level-one nodes, and blue dotted circles denote that of the level-two nodes. The red vertical lines in all the distributions represent the location of the inflection point in level one for the *β* we chose for different interaction networks, *β* = 0.5 for BioGRID. (H) depicts the inflection point *θ* as a function of *β* ranging from 0.05 to 0.95.

